# The Rab GTPase activating protein TBC-2 regulates endosomal localization of DAF-16 FOXO and lifespan

**DOI:** 10.1101/2021.06.04.447106

**Authors:** İçten Meraş, Laëtitia Chotard, Thomas Liontis, Zakaria Ratemi, Benjamin Wiles, Jung Hwa Seo, Jeremy M. Van Raamsdonk, Christian E. Rocheleau

## Abstract

FOXO transcription factors have been shown to regulate longevity in model organisms and are associated with longevity in humans. To gain insight into how FOXO functions to increase lifespan, we examined the subcellular localization of DAF-16 in *C. elegans.* We show that DAF-16 is localized to endosomes and that this endosomal localization is increased by the insulin-IGF signaling (IIS) pathway. Endosomal localization of DAF-16 is modulated by endosomal trafficking proteins. Disruption of the Rab GTPase activating protein TBC-2 increases endosomal localization of DAF-16, while inhibition of TBC-2 targets, RAB-5 or RAB-7 GTPases, decreases endosomal localization of DAF-16. Importantly, the amount of DAF-16 that is localized to endosomes has functional consequences as increasing endosomal localization through mutations in *tbc-2* reduced the lifespan of long-lived *daf-2 IGFR* mutants, depleted their fat stores, and DAF-16 target gene expression. Overall, this work identifies endosomal localization as a mechanism regulating DAF-16 FOXO, which is important for its functions in metabolism and aging.

## INTRODUCTION

Insulin/insulin-like growth factor signaling (IIS) is an evolutionarily conserved pathway that plays an important role in lifespan, development, metabolism, immunity and stress responses from *Caenorhabditis elegans* to humans (Accili and Arden, 2004; Barthel *et al*., 2005; Greer and Brunet, 2005; Murphy and Hu, 2013). IIS-mediated regulation of *C. elegans* FOXO, DAF-16, was first identified via genetic characterization of mutants affecting dauer (an alternative stress resistant larval stage) and life span (Riddle *et al*., 1981; Kenyon *et al*., 1993; Lin *et al*., 1997; Ogg *et al*., 1997). For example, animals that have a mutation in their insulin-like growth factor receptor (IGFR), DAF-2, live almost two times longer than wild type animals in a DAF-16-dependent manner (Kenyon *et al*., 1993). Under favorable conditions, the *C. elegans* DAF-2 IGFR signals through a conserved PI3K/Akt pathway to phosphorylate and inhibit nuclear accumulation of DAF-16 FOXO (Murphy and Hu, 2013). In short, DAF-2 IGFR activation leads to the activation of AGE-1 PI3K which phosphorylates PI(4,5)P_2_ to generate PI(3,4,5)P_3_, which can bind and recruit PDK-1 and AKT-1/2 kinases. DAF-18 PTEN, a lipid and protein phosphatase, countacts AGE-1 PI3K and thus negatively regulates signaling. PDK-1 activates AKT-1/2 which in turn phosphorylates DAF-16 creating binding sites for PAR-5 and FTT-2 14-3-3 scaffold proteins that sequester DAF-16 FOXO in the cytoplasm. During adverse conditions such as starvation, IIS is suppressed and DAF-16 FOXO enters the nucleus to induce the expression of stress response genes. As such, DAF-16 FOXO mediates most *daf-2 IGFR* mutant phenotypes.

Upon activation, the Insulin/IGF receptor is internalized into endosomes, where it can disassociate from its ligand and recycle back to the plasma membrane, or it can be targeted for lysosomal degradation (Bergeron *et al*., 2016). The identification of activated IGFR on endosomes suggested that endosomes can serve as a platform for signaling (Baass *et al*., 1995). Subsequently, several components of the IIS pathway have been shown to localize on endosomes. PTEN localizes on PI(3)P positive endosomes through its C2 domain and has been demonstrated to regulate endosome trafficking via dephosphorylation of Rab7 (Naguib *et al*., 2015; Shinde and Maddika, 2016). Akt2 localizes to Appl-1 and WDFY-2 positive endosomes to get fully activated and regulate Akt2 specific downstream substrates (Schenck *et al*., 2008; Walz *et al*., 2010). 14-3-3 proteins can interact with several Akt phosphorylation targets to regulate their subcellular localization and have been found on endosomes (Jungmichel *et al*., 2014; Marat *et al*., 2017). However, a role for endosome trafficking in regulation of FOXO transcription factors has not been demonstrated to the best of our knowledge.

Rab5 and Rab7 GTPases localize to early and late endosomes respectively and are critical regulators of trafficking to the lysosome, an organelle important for cargo degradation and metabolic signaling (Huotari and Helenius, 2011; Settembre *et al*., 2013). Like many small GTPases, Rabs cycle between a GTP-bound active state and GDP-bound inactive state. This cycling requires guanine nucleotide exchange factors for activation and GTPase Activating Proteins (GAPs) to catalyze GTP hydrolysis and hence Rab inactivation. We previously characterized *C. elegans* TBC-2 as having *in vitro* GAP activity towards RAB-5, and some activity towards RAB-7 (Chotard *et al*., 2010a). Mutations in *tbc-2* result in enlarged late endosomes in several tissues including the intestine, an important site of IIS and metabolic regulation (Libina *et al*., 2003). In addition to early to late endosome maturation, TBC-2 regulates phagosome maturation (Li *et al*., 2009), dense core vesicle maturation (Sasidharan *et al*., 2012) endosome recycling as an effector of RAB-10 and CED-10/Rac (Sun *et al*., 2012; Liu and Grant, 2015) and yolk protein trafficking in oocytes and embryos (Chotard *et al*., 2010b). Yolk protein is prematurely degraded in *tbc-2* mutants. As such, *tbc-2* mutant larvae hatched in the absence of food had reduced survival during L1 (first larval stage) diapause (Chotard *et al*., 2010b).

Here we explore the subcellular localization of DAF-16 and show that DAF-16 localizes on early and late endosomes in *C. elegans* intestinal cells. We found that endosome localization of DAF-16 is regulated by nutrient availability and IIS. We show that endosome localization of DAF-16 is increased through mutations in *tbc-2*, at the expense of nuclear localization. The increased endosomal localization of DAF-16 in *tbc-2* mutants decreases lifespan, fat storage and DAF-16 target gene expression in *daf-2 IGFR* mutant animals. These results demonstrate a role of endosomal localization in the regulation and function of DAF-16 FOXO.

## RESULTS

### TBC-2 and the RAB-5 and RAB-7 GTPases regulate DAF-16 localization to endosomes in the intestine

We previously reported that *tbc-2* is required for survival during L1 diapause (Chotard *et al*., 2010b). Since *daf-16* is also required for survival during L1 diapause (Munoz and Riddle, 2003; Baugh and Sternberg, 2006) we sought to test whether TBC-2 might regulate the nuclear versus cytoplasmic localization of DAF-16. Although we determined that loss of *tbc-2* did not result in precocious development during L1 diapause as seen *daf-16* mutants (Baugh and Sternberg, 2006; Chotard *et al*., 2010b), we found that TBC-2 does in fact regulate DAF-16 localization. Unexpectedly, we found that DAF-16a::GFP (*zIs356*) localized to numerous amorphous vesicles in the intestinal cells of *tbc-2(tm2241)* deletion mutant animals (Figure 1B). Furthermore, we found that DAF-16a::GFP localized to cytoplasmic vesicles in the intestine of wild-type animals (Figure 1A). The percentage of hermaphrodites with DAF-16 vesicles increased during larval development, peaking at L4 and young adults (Figure 1I, Figure 1 - figure supplement 1A). *tbc-2(tm2241)* animals were about twice as likely to have DAF-16 vesicles than wild-type (Figure 1J, Figure 1 - figure supplement 1A). The number of DAF-16 positive vesicles in wild-type can range from zero to hundreds (Figure 1I). Vesicles can be distributed throughout all 20 intestinal cells or be present in high numbers in just a few cells.

**Figure 1.**
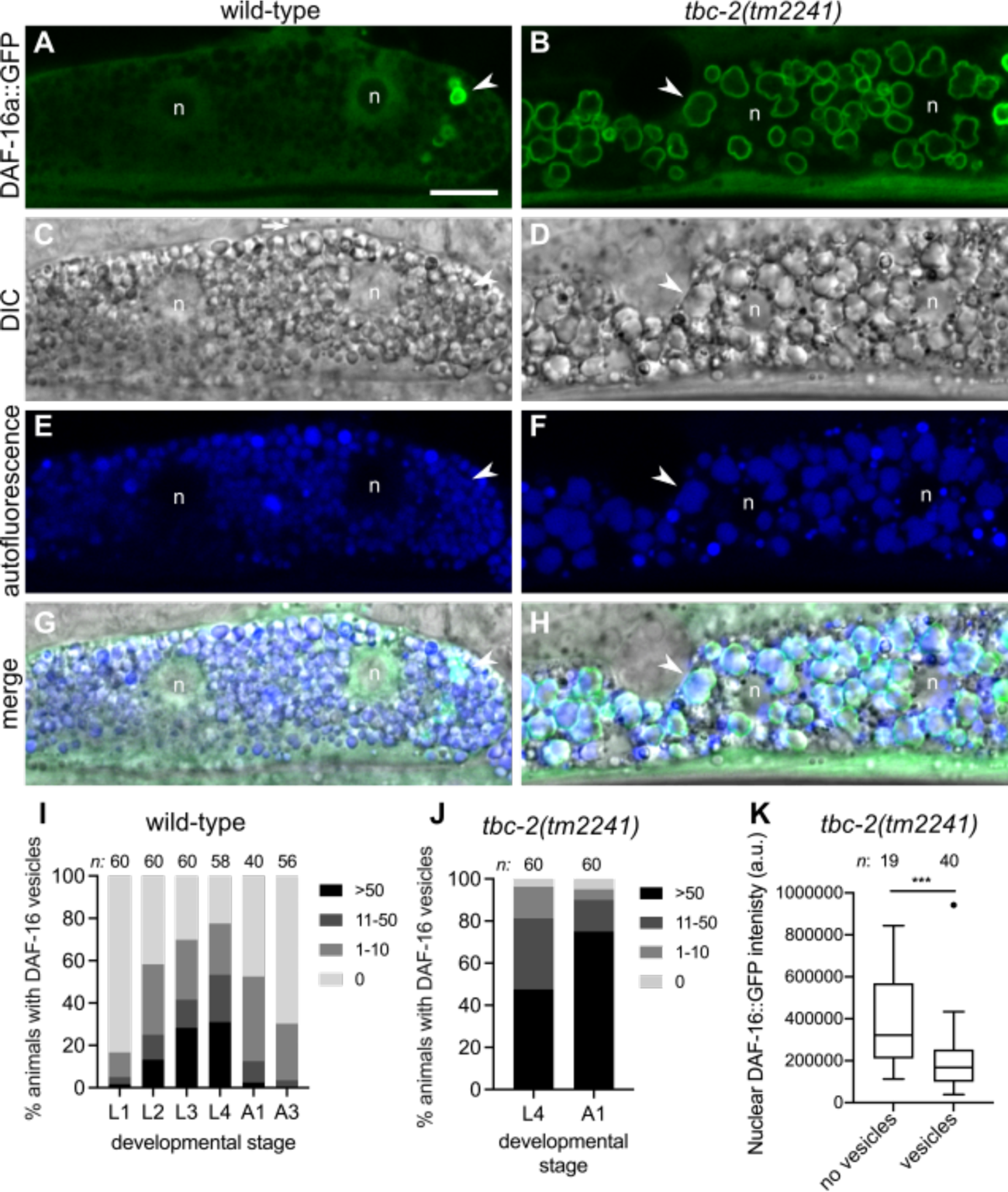
DAF-16 FOXO localizes to vesicles in intestinal cells. (A-H) Representative confocal and differential interference contrast (DIC) images of an intestinal cell of wild-type (A, C, E, G) and *tbc-2(tm2241)* (B, D, F, H) animals expressing DAF-16a::GFP *(zIs356).* DAF-16a::GFP (green) is present on vesicles in both wild-type and *tbc-2(tm2241)* intestinal cells (A and B) that are positive for autofluorescence (blue) present in the the endolysosomal system (E and F). Corresponding DIC (C and D) and merged (G and F) images are shown. A representative vesicle is shown (arrow head) and the two nuclei of the binucleate cell are marked (n). Note that DAF-16a::GFP is excluded from the large nucleoli (A). (I-J) Grouped bar graphs quantifying the percentage of wild-type (I) and *tbc-2(tm2241)* (J) animals with 0, 1-10, 11-50 or >50 DAF-16a::GFP *(zIs356)* positive vesicles at the different larval stages (L1-4) or 1 and 3 day old adults (A1 and A3). (K) A Tukey boxplot of the nuclear intensity in artificial units (a.u.) of DAF-16a::GFP in *tbc-2(tm2241)* intestinal cells with and without DAF-16a::GFP positive vesicles. *n,* total number of animals *** *P<0.001* in an unpaired t test (Prism 8). Scale bar (A), 10μm.

DAF-16a::GFP localization is not sex specific as we found similar numbers of DAF-16 positive vesicles in males as in hermaphrodites (Figure 1 - figure supplement 1B). Although DAF-16a::GFP nuclear localization is low in normal growth conditions, we find that *tbc-2(tm2241)* intestinal nuclei appear to have less nuclear DAF-16a::GFP than wild-type (Figure 1A and B). We quantified the fluorescence intensity of DAF-16a::GFP in nuclei of *tbc-2(tm2241)* intestinal cells comparing nuclei of cells with vesicles versus nuclei of cells without vesicles (Figure 1K). We found that nuclei of cells with vesicles have significantly less nuclear DAF-16a::GFP than cells without vesicles. Thus, DAF-16 localizes to vesicles in wild-type and *tbc-2(tm2241)* animals, and *tbc-2(tm2241)* animals have less nuclear DAF-16a::GFP, likely due to sequestration to cytoplasmic vesicles.

To determine if the localization of the GFP tag or the splice variant or expression levels affected DAF-16 vesicular localization we analyzed three other transgenic strains with lower expression levels; GFP::DAF-16a (*muIs71*), DAF-16a::RFP (*lpIs12*) and DAF-16f::GFP (*lpIs14,* with an alternative N-terminus)(Henderson and Johnson, 2001; Lin *et al*., 2001; Kwon *et al*., 2010). Using the three fluorescent reporters, we detected DAF-16 vesicles in the *tbc-2(tm2241)* mutant, albeit not in the wild-type background (Figure 1 – figure supplement 2G-X).

Overexpression of GFP from *vha-6* intestinal specific promoter, *vhEx1[Pvha-6::GFP]*, did not show significant vesicular localization in wild-type animals (Figure 1 – figure supplement 3A). In *tbc-2(tm2241)* animals GFP showed some vesicular localization and potential aggregates, but much less than seen with DAF-16a::GFP, indicating that the vesicular localization is due to DAF-16 and not the GFP tag. Of note, we found that GFP expression in the *tbc-2* background was visibly stronger than in wild-type (Figure 1 – figure supplement 3B-E), consistent with *vha-6* ranking amongst the top downregulated DAF-16-responsive genes (Tepper *et al*., 2013) and consistent with TBC-2 facilitating DAF-16 nuclear localization.

To determine if endogenous DAF-16 localizes to vesicles we analyzed *daf-16(hq23)*, a DAF-16::GFP line generated by CRISPR/Cas9 genome editing (Zhang *et al*., 2021), and found that endogeneously tagged DAF-16 localized to vesicles in both wild-type and *tbc-2(tm2241)* mutants (Figure 1 – figure supplement 2A-F). To determine if the GFP/RFP tag is driving vesicular localization of DAF-16 we analyzed two *daf-16* alleles endogenously tagged with evolutionarily distant mNeongreen and mKate2 fluorescent proteins (Bhattacharya *et al*., 2019; Aghayeva *et al*., 2020). We found that DAF-16::mNG and DAF-16::mK2 both localized to intestinal vesicles in wild-type animals (Figure 2). Furthermore, the percent wild-type and *tbc-2(tm2241)* animals with endogenous DAF-16::mNG vesicles were comparable to that of the overexpressed DAF-16a::GFP (Figure 2I). Therefore, DAF-16 localization to vesicles is not due to overexpression and unlikely to be an artifact of the fluorescent tag.

**Figure 2.**
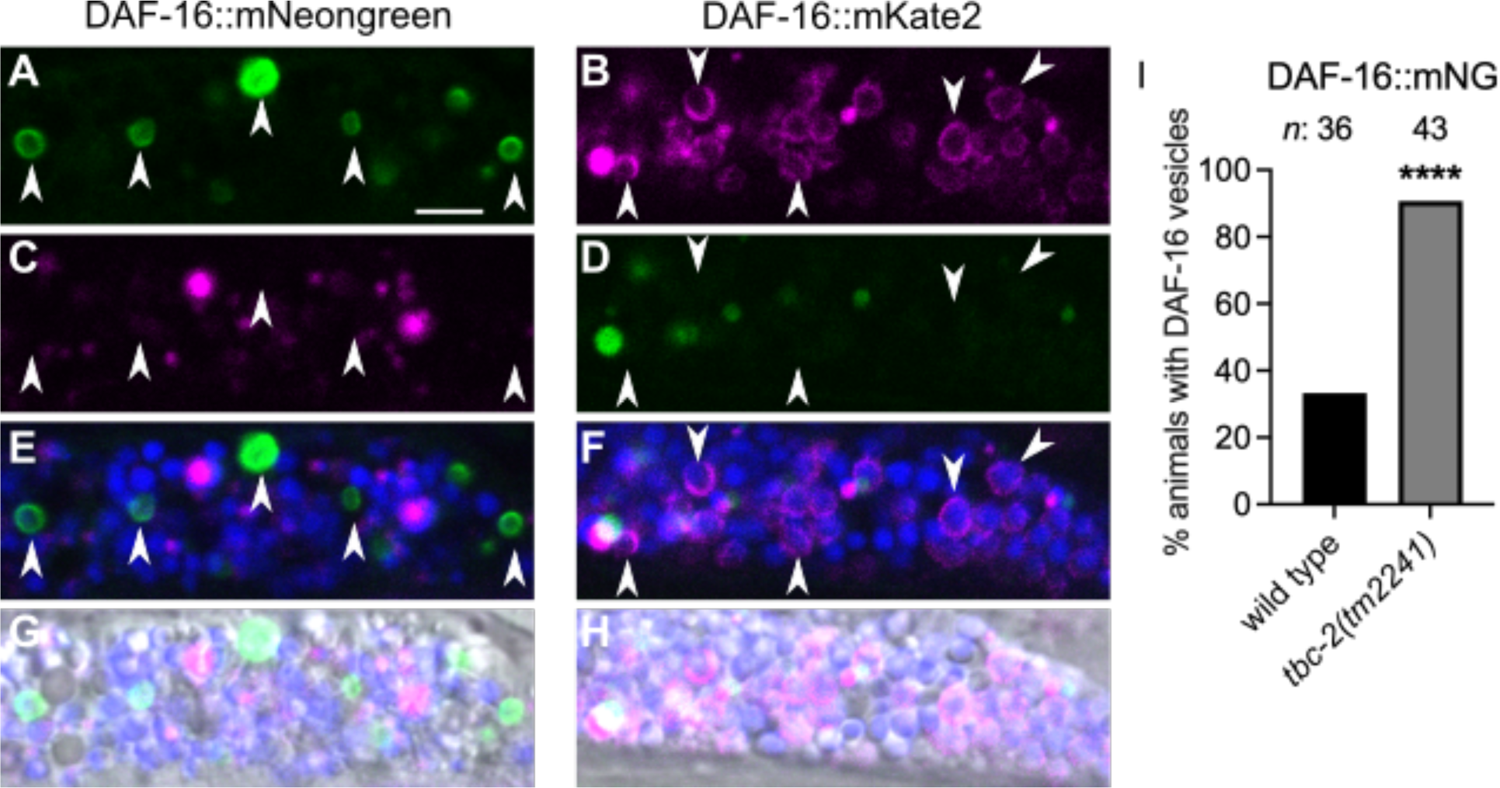
Endogenously tagged DAF-16::mNeongreen and DAF-16::mKate2 localize to vesicles. Representative confocal and differential interference contrast (DIC) images of intestinal cells of *daf-16(ot853[daf-16::linker::mNeongreen::3xFlag::AID])* (A,C,E,G) and *daf-16(ot821[daf-16::mKate2::3xFlag])* (B,D,F,H). Arrows mark DAF-16::mNeongreen positive vesicles in the green channel (A) that are distinct from autofluorescence in the red (C) and blue channels shown as a merge (E). Arrows mark DAF-16::mKate2 positive vesicles in the red channel (B) that are distinct from autofluorescence in the green (D) and blue channels shown as a merge (F). For additional context the fluorescent channels were merged with their corresponding DIC image (G and H). Bar graphs displaying the percent wild-type and *tbc-2(tm2241)* animals with DAF-16::mNG positive vesicles (I). Fisher’s exact test (graphpad.com) was used to determine the statistical difference between conditions. *n*, total number of animals. *\*\*\*\* P<0.0001.* Scale bar (A), 5μm.

The finding that DAF-16 localizes to enlarged vesicles in *tbc-2* mutants and that DAF-16 vesicles contain autofluorescent material (Figure 1E-H, Figure 2E,F), a hallmark of intestinal endolysosomes (Clokey and Jacobson, 1986; Hermann *et al*., 2005; Coburn *et al*., 2013), suggested that DAF-16 vesicles are endosomal. To determine if the DAF-16 vesicles are endosomes, we co-expressed DAF-16a::GFP with RFP::RAB-5, an early endosomal marker, or with mCherry::RAB-7, a late endosomal marker (Chavrier *et al*., 1990; Sato *et al*., 2005; Chen *et al*., 2006). We found that in wild-type animals, DAF-16a::GFP localizes to a subset of RAB-5 and RAB-7-positive endosomes (Figure 3A-F). Thirty percent of DAF-16a::GFP vesicles were RAB-5 positive (n=959) while 5% were RAB-7 positive (n=365). Furthermore, we tested the genetic requirements for *rab-5* and *rab-7* for DAF-16 localization by RNAi. We found that *rab-5(RNAi)* and *rab-7(RNAi)* knockdown significantly decreased the number of wild-type and *tbc-2(tm2241)* animals with DAF-16a::GFP vesicles (Figure 3G and H). Together, these data are consistent with DAF-16 localizing to a sub-population of endosomes or endosome-like vesicles in the intestinal cells. It further implicates endosomal regulators TBC-2, RAB-5 and RAB-7 as regulators of DAF-16.

**Figure 3.**
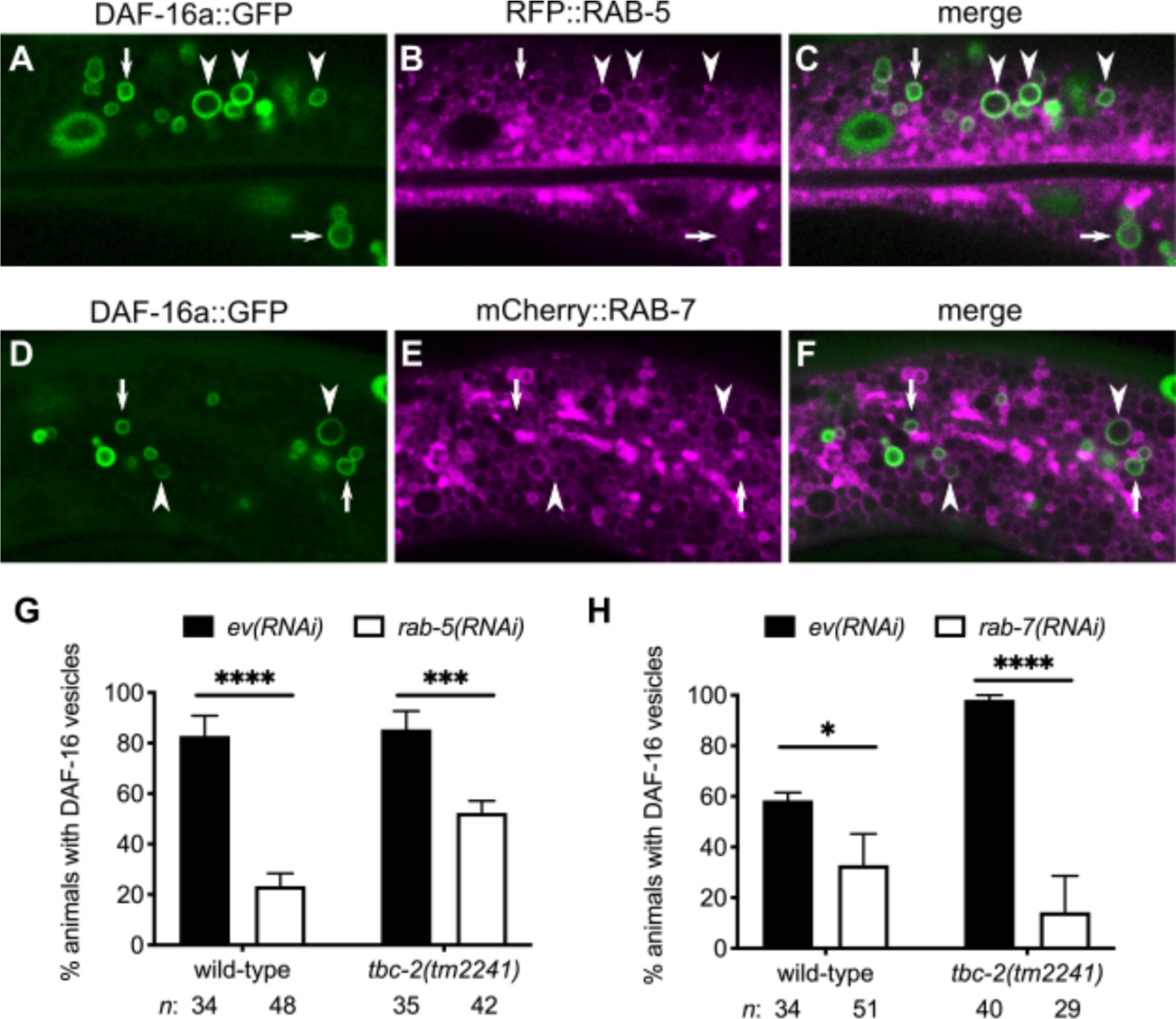
RAB-5 and RAB-7 GTPases promote DAF-16 FOXO localization to endosomes. (A-F) Representative confocal images of the intestinal cells of animals expressing DAF-16a::GFP (*zIs356)* (green) and either RFP::RAB-5 (*pwIs480*) (A-C) or mCherry::RAB-7 (*pwIs429*) (magenta) (D-F). Arrowheads mark examples of vesicles positive for both DAF-16a::GFP and either RFP::RAB-5 or mCherry::RAB-7. Arrows mark examples of DAF-16a::GFP vesicles that are not positive for either RFP::RAB-5 or mCherry::RAB-7. (G and H) Bar graphs displaying the mean (and SEM) of the percent wild-type and *tbc-2(tm2241)* animals with DAF-16a::GFP (*zIs356*) vesicles fed bacteria expressing control empty vector (ev) RNAi (black bars) compared to animals fed *rab-5(RNAi)* or *rab-7(RNAi)* (white bars) from three independent experiments. Fisher’s exact test (graphpad.com) was used to determine the statistical difference between conditions. *n*, total number of animals. * *P<0.05, *** P<0.001, **** P<0.0001*.

### Acute starvation suppresses DAF-16 localization to endosomes

Nuclear localization of DAF-16 is modulated by nutrient availability. As such, starvation promotes DAF-16 cytoplasmic-to-nuclear shuttling (Henderson and Johnson, 2001). To determine if endosomal DAF-16 can translocate to the nucleus, we tested the effect of acute starvation on DAF-16a::GFP localization to endosomes. We found that starvation strongly suppressed the localization of DAF-16 to endosomes in both wild-type and *tbc-2(tm2241)* mutants (Figure 4A; Figure 4 – figure supplement 1). Upon re-feeding, DAF-16 relocalized to endosomes after 1-2 hours, in both wild-type and *tbc-2(tm2241)* mutants. Thus, DAF-16 localization on endosomal membranes is regulated by nutrient availability.

**Figure 4.**
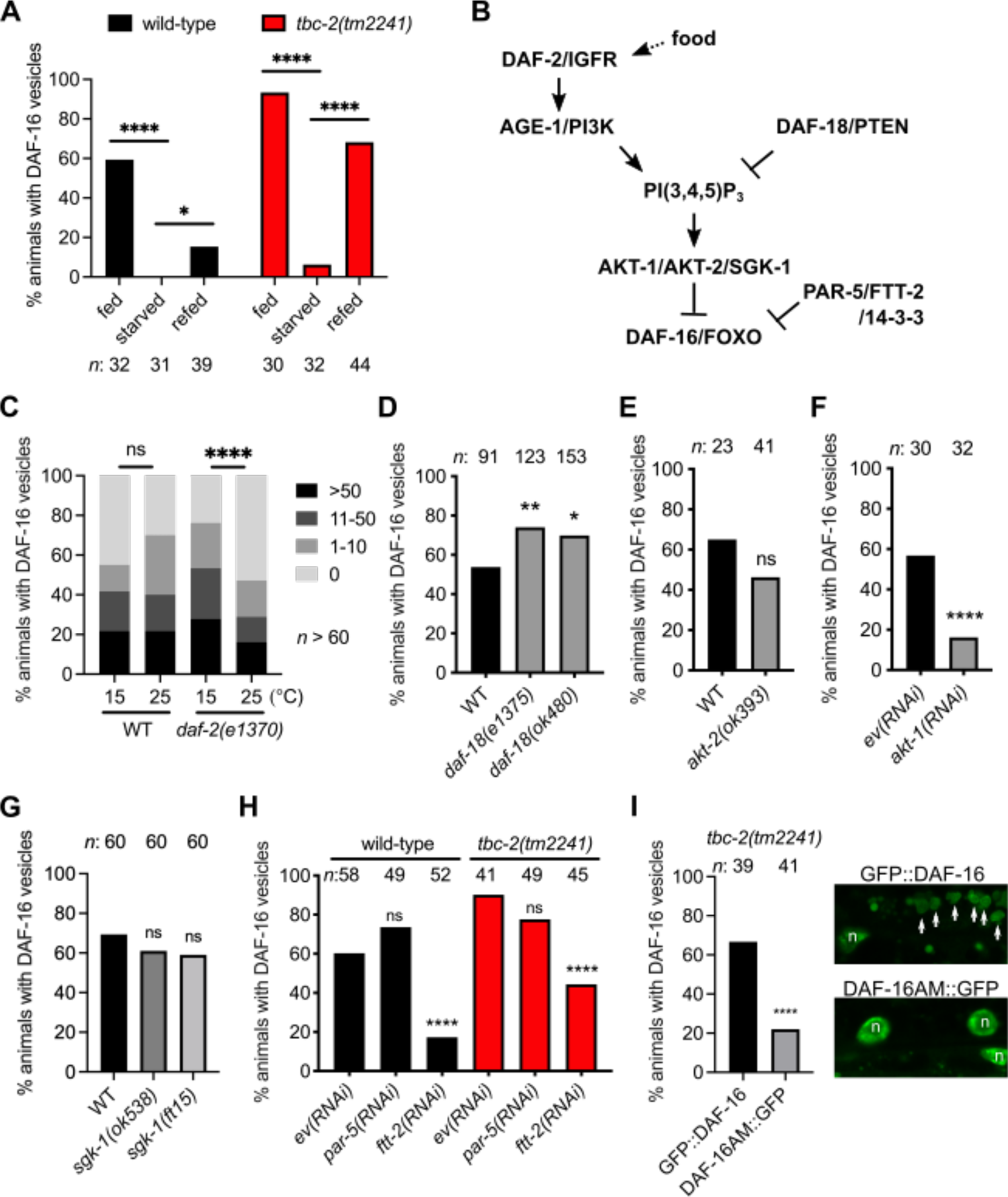
DAF-16 FOXO endosome localization is regulated by IIS and nutrient availability. (A) Bar graphs of the percent wild-type (black) and *tbc-2(tm2241)* (red) animals with DAF-16a::GFP (*zIs356*) vesicles in fed animals, animals starved between 4 and 5 hours and starved animals that have been refed for 1 to 2 hours. (B) Diagram of IIS-mediate regulation of DAF-16/FOXO. (C) Grouped bar graphs quantifying the percentage of wild-type and *daf-2(e1370)* L4 larvae (cumulative data from 3 independent strains) with 0, 1-10, 11-50 or >50 DAF-16a::GFP *(zIs356)* positive vesicles at 15°C or shifted overnight to 25°C. (D) Bar graphs of percent wild-type and *daf-18(e1375)* (cumulative data from 3 independent strains) and *daf-18(ok480)* (cumulative data from 5 independent strains) with DAF-16a::GFP (*zIs356*) vesicles. (E) Bar graphs of percent wild-type and *akt-2(ok393)* (cumulative data from 2 independent strains) with DAF-16a::GFP (*zIs356*) vesicles. (F) Bar graphs of percent wild-type animals treated with control empty vector RNAi and *akt-1(RNAi)* with DAF-16a::GFP (*zIs356*) vesicles. (G) Bar graphs of percent wild-type, *sgk-1(ok538)* and *sgk-1(ft15)* with DAF-16a::GFP (*zIs356*) vesicles. (H) Bar graphs of the percent wild-type (black) and *tbc-2(tm2241)* (red) animals fed control empty vector RNAi, *par-5(RNAi)* and *ftt-2(RNAi)* with DAF-16a::GFP (*zIs356*) vesicles. (I) Bar graph of the precent *tbc-2(tm2241)* animals with GFP::DAF-16a (*muIs71*) or DAF-16a^AM^::GFP *(muIs113)-*positive vesicles. Representative images of GFP::DAF-16a (*muIs71*) (top) and DAF-16a^AM^::GFP *(muIs113)* (bottom) are shown. GFP::DAF-16a vesicles (arrows) and the nuclei (n). Fisher’s exact test (graphpad.com) was used to determine the statistical difference between conditions. *n*, total number of animals. ns, not significant, * *P<0.05, ** P<0.01, *** P<0.001, **** P<0.0001*.

### Insulin/IGF signaling regulates DAF-16 localization to endosomes

To determine if IIS regulates DAF-16 localization to endosomes we analyzed the effect of disrupting IIS on DAF-16a::GFP localization (see Figure 4B for reference). Firstly, we analyzed a hypomorphic mutant of the insulin/IGF receptor, *daf-2(e1370)*, in which IIS is reduced, particularly at higher temperatures (Swanson and Riddle, 1981). We compared DAF-16a::GFP localization in three independent *daf-2(e1370)* strains at 15°C and shifted overnight to 25°C to enhance disruption of DAF-2. All three showed a significant reduction in the number of DAF-16 positive vesicles at 25°C, while the wild-type strain did not (Figure 3C), indicating that DAF-2 promotes DAF-16 localization to endosomes.

DAF-18, the homolog of human tumor suppressor protein PTEN, acts as a negative regulator of the IIS pathway counteracting AGE-1 PI3K signaling by dephosphorylating the 3’phosphate on the PI(3,4,5)P _3_ converting it to PI(4,5)P_2_ (Ogg and Ruvkun, 1998). We analyzed DAF-16a::GFP localization in the *daf-18* reference allele, *e1375.* We generated three independent *daf-18(e1375); zIs356* strains, but only two strains had increased endosome localization of DAF-16. However, combined data from the three strains had a statistically significant increase in the number of animals with DAF-16 endosomes (Figure 4D). Since *daf-18(e1375)* is possibly a non-null allele, we analyzed a deletion allele, *daf-18(ok480).* We found that five independent *daf-18(ok480); zIs356* strains had more DAF-16 endosomes than wild-type that when combined was statistically significant (Figure 4D). Since DAF-16 endosome localization is mildly increased in the background of two distinct *daf-18* alleles, it is consistent with increased IIS causing an increase in DAF-16 localization to endosomes.

*C. elegans* AKT-1 and AKT-2 function upstream of, and can phosphorylate, DAF-16 (Paradis and Ruvkun, 1998; Hertweck *et al*., 2004). The SGK-1 Serum Glucorticoid Kinase homolog interacts with AKT kinases and has been shown to regulate DAF-16 nuclear localization (Hertweck *et al*., 2004). We tested which of these kinases might regulate DAF-16 localization to endosomes. We found that animals with an *akt-2(ok393)* deletion mutation, there was an insignificant decrease in DAF-16 endosomal localization in two independent strains (Figure 4E). However, in *akt-1*(*RNAi)* animals, we observed a significant decrease in DAF-16 endosomal localization where DAF-16 localized mainly to the nucleus, compared to the control animals in three independent experiments (Figure 4F). Neither an *sgk-1(ok538)* deletion nor an *sgk-1(ft15)* gain-of-function allele affected DAF-16 localization to endosomes (Hertweck *et al*., 2004; Jones *et al*., 2009)(Figure 4G). Therefore, AKT-1 is the main kinase regulating DAF-16 localization to endosomes.

Phosphorylation of FOXO proteins by Akt creates binding sites for 14-3-3 proteins, which promote cytoplasmic retention of FOXO (Brunet *et al*., 1999). *C. elegans* has two genes that encode for 14-3-3 proteins that interact with DAF-16, *par-5* and *ftt-2* (Wang and Shakes, 1997; Morton *et al*., 2002; Wang *et al*., 2006). We found that *ftt-2(RNAi)*, but not *par-5(RNAi)*, suppressed the localization of DAF-16 to endosomes (Figure 4H), suggesting that FTT-2 regulates DAF-16 endosomal localization.

To determine if AKT phosphorylation regulates DAF-16 localization to endosomes we tested if DAF-16a^AM^::GFP *muIs113*, in which the four consensus AKT phosphorylation sites (T54A, S240A, T242 and S314A) are mutated to alanine (Lin *et al*., 2001), could still localize to vesicles in *tbc-2(tm2241)* mutants. Since the expression levels are lower than DAF-16a::GFP *zIs356*, we used GFP::DAF-16a *muIs71* with comparable expression levels as control (Figure 4 – figure supplement 2). We found that the number of *tbc-2(tm2241)* animals with DAF-16a^AM^::GFP positive vesicles was significantly less than the GFP::DAF-16a control (Figure 4I). Together, these data demonstrate that IIS regulates endosomal DAF-16 through AKT-1 specific phosphorylation and FTT-2 14-3-3.

### TBC-2 is required for lifespan extension and increased fat storage of *daf-2(e1370)* mutants

To determine if the increased endosomal localization of DAF-16 seen in *tbc-2* mutants affects DAF-16 activity, we tested if TBC-2 regulates adult lifespan. We found that the *tbc-2(sv41)* and *tbc-2(tm2241)* deletion alleles had similar lifespans as compared to wild-type (Figure 5A and B; Figure 5 – figure supplement 1). As expected, *daf-2(e1370)* animals lived significantly longer than wild-type. We found that both *tbc-2* alleles significantly shortened the lifespan of *daf-2(e1370)* animals (Figure 5A and B; Figure 5 – figure supplement 1). Thus, TBC-2 is not required for normal lifespan, but is partly required for the extended lifespan of *daf-2(e1370)* mutants.

**Figure 5.**
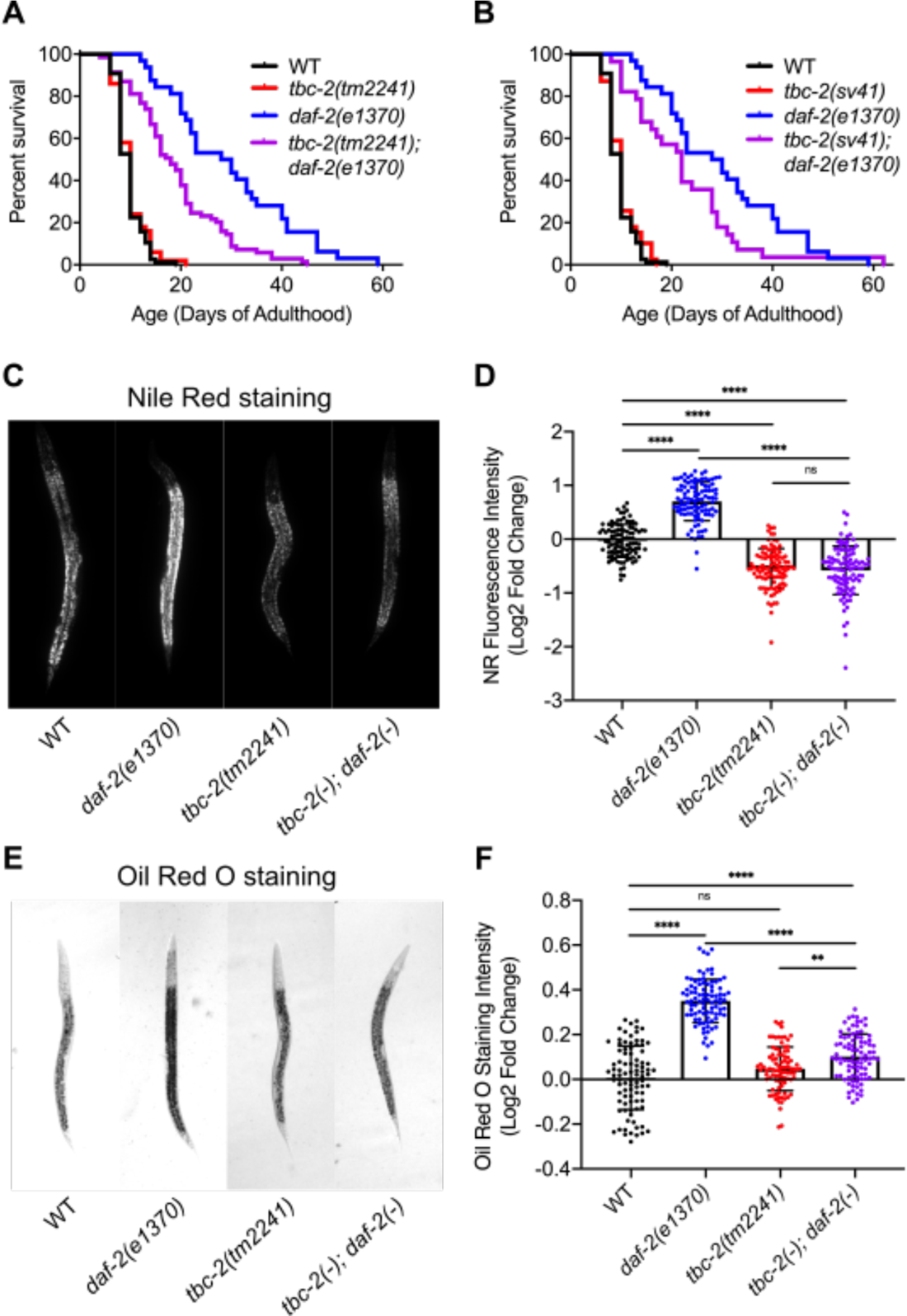
TBC-2 is required for full lifespan extension and increased fat storage resulting from decreased insulin/IGF signaling. (A,B) Percent survival curve of adult animals of wild-type and *daf-2(e1370)* with either *tbc-2(tm2241)* and *tbc-2(tm2241); daf-2(e1370)* (A) or *tbc-2(sv41)* and *tbc-2(sv41); daf-2(e1370)* (B) genotypes. The survival of all strains are statistically different from *daf-2(e1370)* animals as determined by a Mantel-Cox log-rank test: *tbc-2(tm2241); daf-2(e1370)* (*P<0.001*) and *tbc-2(sv41); daf-2(e1370)* (*P=0.0314*). Cumulative data from three independent replicates of 50 young adults for a total of 150. (C,D) Log2 fold change in Nile Red (NR) fluorescence intensity (C) or Oil Red O staining intensity (D) comparing wild-type, *daf-2(e1370)*, *tbc-2(tm2241)* and *tbc-2(tm2241); daf-2(e1370)* fixed L4 hermaphrodites. Statistical analysis was done using a student’s t-test with a one way analysis of variance (ANOVA). ns, not significant. ** *P* <0.01, **** *P* <0.0001.

To assess TBC-2’s contribution to other *daf-2* mutant phenotypes, we tested whether TBC-2 is required for the increased fat storage of *daf-2(e1370)* mutants. Consistent with previous findings we found that *daf-2(e1370)* had significantly more fat than wild-type animals as determined by Nile Red and Oil Red O staining of fixed L4 larvae (Kimura *et al*., 1997; O’Rourke *et al*., 2009) (Figure 5C-F). We found that *tbc-2(tm2241)* larvae had less fat than wild-type with Nile Red staining, but not with Oil Red O staining (Figure 5D and F). This difference might reflect the lower sensitivity of Oil Red O for quantifying lipid abundance as compared to Nile Red (Escorcia *et al*., 2018). Interestingly, we found that *tbc-2(tm2241)* significantly suppressed the *daf-2(e1370)* increased lipid staining by both Nile Red and Oil Red O (Figure 5C-F). Thus, TBC-2 is required for the increased fat storage of *daf-2(e1370)* mutants.

The fact that TBC-2 is required for the increased longevity and increased fat storage of *daf-2(e1370)* mutants suggests that TBC-2 could be a negative regulator of IIS. Therefore, we tested if *tbc-2* mutants regulate DAF-16 target gene expression in *daf-2(e1370)* animals. We used qRT-PCR to measure the expression of DAF-16 target genes in wild-type, *daf-2(e1370), tbc-2* mutants and *tbc-2; daf-2* double mutants. Consistent with previous reports, the expression of six of the DAF-16 target genes were upregulated in *daf-2(e1370)* animals (Figure 6) (Honda and Honda, 1999; Murphy *et al*., 2003; Ackerman and Gems, 2012; Tepper *et al*., 2013). While *tbc-2(tm2241)* and *tbc-2(sv41)* did not appreciably alter DAF-16 target gene expression relative to wild-type, both reduced the expression of *sod-3, dod-3, gpd-2* and *icl-1* in *daf-2(e1370)* mutants, while *mtl-1* and *ftn-1* were not significantly decreased (Figure 6). The fact that TBC-2 was required for the increased expression of four of six DAF-16 target genes with elevated expression in *daf-2(e1370)* mutants suggests a more specific role for TBC-2 in regulating DAF-2 to DAF-16 signaling.

**Figure 6.**
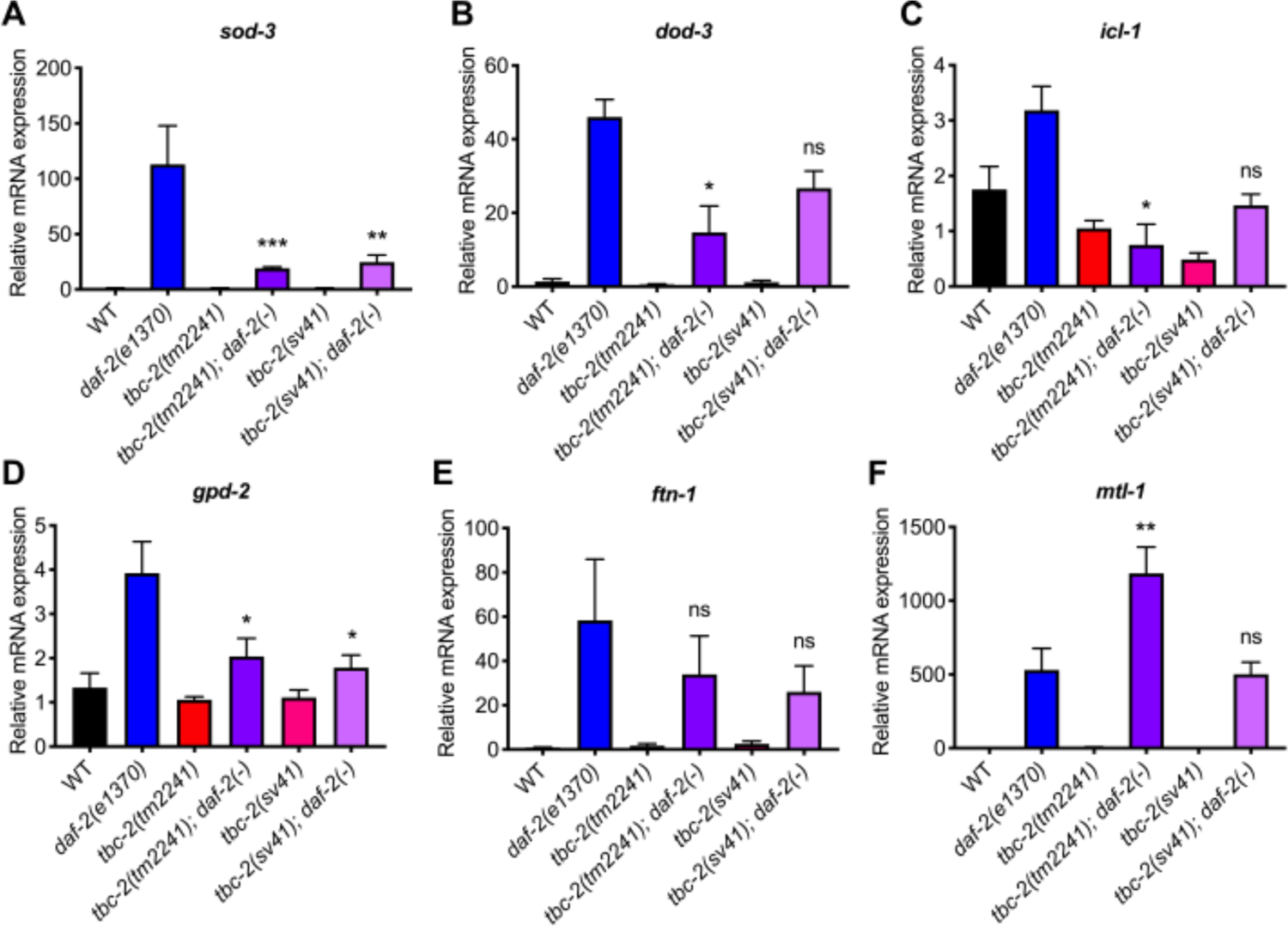
TBC-2 is required for the increased DAF-16 target gene expression resulting from decreased insulin/IGF signaling Quantitative RT-PCR analysis of DAF-16 target gene expression in wild-type, *daf-2(e1370)*, *tbc-2(tm2241), tbc-2(tm2241); daf-2(e1370), tbc-2(sv41),* and *tbc-2(sv41); daf-2(e1370).* Statistical analysis was performed using an Ordinary one-way ANOVA – Tukey’s multiple comparisons test. Shown are comparisons of *tbc-2(tm2241); daf-2(e1370)* and *tbc-2(sv41); daf-2(e1370)* versus *daf-2(e1370)* single mutants. ns, not significant. *p<0.05, **p<0.01, ***p<0.001.

## DISCUSSION

Endosome trafficking and signal transduction are intimately linked processes regulating signal propagation, specificity and attenuation (Miaczynska, 2013; Bergeron *et al*., 2016). However, there remains a large knowledge gap in the spatial regulation of cell signaling and where downstream transcription factors are regulated. We identified a previously unknown localization for the DAF-16 FOXO transcription factor on endosomes in *C. elegans*. Endosome localization is limited by the TBC-2 Rab GAP. Loss of *C. elegans* TBC-2 results in increased endosomal localization of DAF-16 at the expense of nuclear localization. As such, *C. elegans* TBC-2 is partly required for several *daf-2 IGFR* mutant phenotypes including lifespan extension, increased fat storage, and increased DAF-16 target gene expression that result from DAF-16 nuclear translocation. DAF-16 endosome localization is largely dependent on IIS consistent with this being a phosphorylated, inactive pool of DAF-16. Together our data show a role for the TBC-2 Rab GAP in regulating the balance of nuclear versus endosomal localization of the DAF-16 transcription factor.

### DAF-16 localizes to endosomes

We were surprised to find that DAF-16 FOXO localizes to a subset of RAB-5 and RAB-7 endosomes in wild-type animals. Many studies have used DAF-16 cytoplasmic versus nuclear localization to assess IIS activity under various conditions. We assume that DAF-16 positive endosomes were not discovered earlier as nuclear translocation can be assessed at low magnification where DAF-16 positive endosomes are not apparent, and DAF-16 positive endosomes are not present in every cell or every animal. Furthermore, we first took notice of DAF-16 positive endosomes in the *tbc-2* mutant background where they are more prominent. These are likely not an artefactual consequence of overexpression, as we see these vesicles in the endogenously tagged *daf-16::GFP* and we do not see similar vesicles in a GFP overexpression strain. DAF-16::mNG and DAF-16::mK2 also localize to vesicles indicating that membrane localization is unlikely to be an artifact of the GFP tag. Additionally, DAF-16-vesicles are regulated by IIS.

Many components of the IIS pathway localize to endosomes in mammalian cells including active insulin receptor and downstream signaling components such as PI3K, Akt, PTEN and 14-3-3 proteins (Khan *et al*., 1986; Christoforidis *et al*., 1999; Balbis *et al*., 2000; Braccini *et al*., 2015; Naguib *et al*., 2015; Bergeron *et al*., 2016; Marat *et al*., 2017; Kim *et al*., 2018). In the case of Akt and PTEN, both have demonstrated roles in regulation of endosome trafficking independent of IIS (Shinde and Maddika, 2016, 2017; Mondin *et al*., 2019; Hsu *et al*., 2020). On the other hand, the PI3Ks are Rab5 effectors and Rab5 has been shown to promote Akt activity on endosomes (Christoforidis *et al*., 1999; Shin *et al*., 2005; Su *et al*., 2006; Cheng *et al*., 2009; Braccini *et al*., 2015). While endosomal localization of FOXO proteins has not been reported to the best of our knowledge, knockdown of Rab5 in mouse liver results in a strong increase in phosphorylated FOXO1 (Zeigerer *et al*., 2015). This is contrary to the finding that Rab5 promotes Akt phosphorylation (Su *et al*., 2006; Braccini *et al*., 2015), which could be a consequence of indirect regulation or suggest tissue-specific regulation. The fact that TBC-2 is a RAB-5 GAP is consistent with increased RAB-5 activity promoting DAF-16 localization on endosomes. This is further supported by the fact that *rab-5* and *rab-7* RNAi knockdown reduces the number of animals with DAF-16 vesicles in both wild-type and *tbc-*2 mutants. Given the importance of RAB-5 for endosome trafficking, it is difficult to parse whether RAB-5 is promoting a platform for DAF-16 localization or if it also has a role in IIS.

We demonstrated that DAF-16 localized to a subset of RAB-5 and RAB-7 positive endosomes. Since RAB-5 and RAB-7 promote trafficking to the lysosome and promote receptor tyrosine kinase degradation, it is possible that DAF-16-positive endosomes are signaling endosomes, in which case we would expect other upstream signaling components might be present. Consistent with that hypothesis, knockdown of IIS reduces the number of animals with DAF-16 vesicles. However, when analyzing DAF-16 localization in *daf-2(e1370)* mutants at 15°C vs. 25°C, we find that while there is a reduction in the number of DAF-16 endosomes, these endosomes are noticeably fainter at 25°C. This suggests that there are not necessarily less endosomes being generated, but rather less DAF-16 on the vesicles which may be inconsistent with these being signaling endosomes derived from DAF-2 internalization at the plasma membrane. On the other hand, the fact that Akt and 14-3-3 can localize to endosomes in mammalian cells (Balbis *et al*., 2000; Braccini *et al*., 2015; Marat *et al*., 2017) and that AKT-1 and FTT-2 promote DAF-16 localization to endosomes, suggests that these proteins might recruit DAF-16 onto endosomes rather than DAF-16 interacting directly with membranes. Future studies should test whether DAF-2 IGFR and downstream IIS components actively recruit DAF-16 to endosomes or whether IIS has a passive role. IIS inhibition of nuclear DAF-16 could result in increased DAF-16 in the cytoplasm where it can bind endosomes.

If these are not signaling endosomes, then what are the DAF-16 endosomes? Since RAB-5 and RAB-7 are also regulators of autophagy, so it is possible that these endosomes contribute to degradation of inactive excess DAF-16. There is precedent for selective autophagy in the degradation of the GATA4 transcription factor (Kang *et al*., 2015). Alternatively, these endosomes could serve as a reservoir of inactive DAF-16 that can be quickly mobilized if environmental stress is encountered. For example, we found that acute starvation is a potent regulator of DAF-16 endosome localization, even in the *tbc-2* mutant background.

We find it interesting that there is such variability within the population, and amongst the intestinal cells in a given animal, as to whether there will be DAF-16 positive endosomes or not. It suggests that each intestinal cell autonomously senses changes in IIS, or possibly other nutrient and stress sensing pathways, to regulate DAF-16 localization. Then, why is endosomal DAF-16 more prominent in *tbc-2* mutants? One explaination would be that the expansion of endosomal membranes in a *tbc-2* mutant create more storage space for inactive DAF-16. Another would be that IIS or other pathways are more active in *tbc-2* mutants, or some combination of the two.

GLP-1/Notch signaling in the germline, AMPK, JNK and LET-363/mTor signaling additionally regulate DAF-16 and could be subject to regulation by TBC-2, particularly mTor which localizes to lysosomes (Oh *et al*., 2005; Berman and Kenyon, 2006; Greer *et al*., 2007; Sancak *et al*., 2008; Robida-Stubbs *et al*., 2012; Sun *et al*., 2017).

### A direct role for TBC-2 in regulating *daf-2(e1370) IGFR* mutant phenotypes

Our finding that *tbc-2* was required for the extended lifespan and increased fat storage of *daf-2(e1370)* mutants suggests that TBC-2 might have a more specific role related to IIS. However, TBC-2 is not the first endosomal regulator required for lifespan extension of *daf-2(e1370)* mutants. *C. elegans* BEC-1, a homolog of human Beclin1, is a regulator of autophagy and endosome trafficking, and *bec-1* mutants accumulate large late endosomes in the intestinal cells (Melendez *et al*., 2003; Ruck *et al*., 2011; Law *et al*., 2017). Mutations in *bec-1* suppress the increased lifespan of *daf-2(e1370)*, and were reported to be required for the increased fat storage (Melendez *et al*., 2003). Similarly, other autophagy regulators, *atg-7* and *atg-12*, have been shown to be required for *daf-2* longevity (Hars *et al*., 2007). An RNAi screen identified regulators of endosome to lysosome trafficking, including RAB-7, and components of the ESCRT and HOPS complexes as being required for the lifespan extension phenotypes of *daf-2 IGFR* mutants (Samuelson *et al*., 2007). However, the mechanisms by which they regulate lifespan are not known. In *daf-2(e1370)* mutants, there is an increase in autophagy and lysosome function, both of which are required for the extended lifespan (Melendez *et al*., 2003; Lapierre *et al*., 2013; Guo *et al*., 2014; Sun *et al*., 2020), consistent with increased stress resistance contributing to increased longevity. Since the human homologs of TBC-2 are implicated in autophagy (Behrends *et al*., 2010; Popovic *et al*., 2012; Carroll *et al*., 2013; Toyofuku *et al*., 2015), and *tbc-2* mutants accumulate autophagy protein LGG-1 (LC3/Atg8) in enlarged endosomes (Chotard *et al*., 2010a), it is possible that TBC-2 also regulates autophagy and thus *daf-2(-)* longevity. However, our findings that TBC-2 regulates DAF-16 nuclear vs. endosome localization and that TBC-2 is required for the DAF-16 target gene expression in *daf-2* mutants demonstrate that TBC-2 has a more direct role in IIS, as opposed to being required for downstream cellular responses. It will be interesting to test if TBC-2 regulates DAF-16-independent mechanisms of longevity as well as to determine if other endosome and autophagy-regulating genes can regulate DAF-16 localization to endosomes.

In conclusion, we demonstrate that the DAF-16 FOXO transcription factor localizes to endosomes. This endosomal localization of DAF-16 FOXO is regulated by both IIS and the TBC-2 family of Rab5 and Rab7 GAPs. TBC-2 promotes the nuclear localization of DAF-16 FOXO, and this localization has functional effects on both longevity and metabolism through modulation of DAF-16 target gene expression. Our data suggest that endosomes serve as an important location for DAF-16 FOXO transcription factor regulation, and suggest that endomembranes may function as a site of transcription factor regulation.

## METHODS

### *C. elegans* genetics and strain construction

*C. elegans* strains were cultured as described in Wormbook (www.wormbook.org). The *C. elegans* N2 Bristol strain was the wild-type parent strain and HB101 *E. coli* strain was used as a food source. Both were obtained from the Caenorhabditis Genetic Center (CGC) as were many of the strains used in this study (Key Resource Table). New strains were constructed using standard methods and the presence of mutations were confirmed by PCR and DNA sequencing.

### RNAi experiments

*C. elegans* RNAi feeding experiments were conducted essentially as described in (Kamath *et al*., 2001). RNAi feeding clones were obtained from the Ahringer RNAi library and confirmed by sequencing (Key resources table)(Fraser *et al*., 2000; Kamath *et al*., 2003). The L4440 empty RNAi feeding vector transformed into HT115(DE3) was used as a negative control (Timmons and Fire, 1998).

### Microscopy

DAF-16 vesicular localization was analyzed in the intestinal cells of hermaphrodites. Hermaphrodite worms at the L4 stage were imaged alive at room temperature unless it is stated otherwise. Animals were picked and mounted onto 4% agarose pads, animals were anesthetized with levamisole.

Differential interference contrast (DIC) and fluorescent imaging were performed with an Axio Imager A1 compound microscope with a 100×1.3 NA Plan-Neofluar oil-immersion objective lens (Zeiss) and images were captured by using an Axio Cam MRm camera and AxioVision software (Zeiss). Confocal microscopy was performed on an Axio Observer Z1 LSM780 laser scanning confocal microscopes with a 63×1.4 NA Plan-Apochromat oil-immersion objective lens (Zeiss) in a multi-track mode using an argon multiline laser (405 nm excitation for autofluoresence, 488 nm excitation for GFP and a 561/ 594 nm excitation for mCherry/RFP). Images were captured with a 32 channel GaAsP detector and ZEN2010 image software. Raw data was analyzed using Fiji (ImageJ) or Zen 2010 Lit programs, and images were modified by using Fiji (ImageJ).

To compare DAF-16 (*zIs356*) nuclear intensity in cells with or without DAF-16 vesicles, animals at L4 stage were imaged using an LSM780 scanning laser microscope. To ensure consistency, only the anterior most intestine cells were imaged. First, the nucleus of intestinal cells were focused under bright field and without changing the position, GFP, autofluoresence and bright field signals were imaged. Each animal was imaged using the same confocal settings. After data collection, each nucleus was categorized as a nucleus with adjacent DAF-16-positive vesicles or a nucleus without any DAF-16 positive vesicles. Total GFP intensity inside the nucleus was measured using Fiji (Image J) software. The nucleus is circled using DIC/Bright Field and autofluorescence channels as reference. Since intestinal cells have two nuclei per cells, if there are two nuclei within the focus, their GFP intensity is averaged for statistical analysis. Prism 8 (GraphPad) were used to graph the data and determine statistical analysis using an unpaired t-test.

### Starvation-Refeeding and Temperature Shift Experiments

For starvation and refeeding experiments animals were synchronized at the L1 stage and grown NGM plates with HB101 *E. coli* till the L4 stage. Then animals were collected washed 3 times for 5 mins with M9 buffer to remove bacteria in their gut. After the third wash animals were plated to regular NGM plates with or without HB101 *E. coli*, and incubated for 4-5 hours before scoring. Animals are scored for the presence of DAF-16::GFP-positive vesicles the intestine using an A1 Zeiss microscope. After 4h-5h of starvation, animals were harvested with M9 buffer from the starved plates and washed once with M9 buffer and plated to NGM plates with HB101 *E. coli* and incubated for 1-2 hours at 20°C before scoring.

### Life span analysis

Replicate strains were maintained for several generations prior to beginning the lifespan assays which were conducted at 20°C. For each strain 25 young adult hermaphrodites were picked to two NGM plates without FUDR seeded with HB101 *E. coli* for three independent replicates totaling 150 animals. Strains were coded and scored blindly to reduce bias. Animals were transferred to fresh NGM plates to avoid contamination and getting crowded out by their progeny. Animals were scored every 2-3 days and were considered dead when they stop exhibiting spontaneous movement and fail to move in response to 1) a gentle touch of the tail, 2) a gentle touch of the head, and 3) gently lifting the head. Animals that die of unnatural causes (internal hatching of embryos, bursting, or crawling off the plate) are omitted. Graphs and statistics were done using Graphpad Prism. None of the strains used in the lifespan assays carry the *fln-2(ot611)* mutation found in a N2 male stock strain and found to extend median lifespan (Gems and Riddle, 2000; Zhao *et al*., 2019).

### Fat staining

L4 animals were fixed for staining with Nile Red (Invitrogen) or Oil Red O (Sigma-Aldrich) as previously described (Escorcia *et al*., 2018). Imaging and analysis was done as previously described (Ratemi *et al*., 2019). Graphs and statistics were done using Graphpad Prism.

### Quantitative real-time RT-PCR

*C. elegans* RNA was isolated from young adults maintained at 15°C using TRIZOL reagent (Invitrogen). 1 ug of RNA was reverse transcribed into cDNA using the High-Capacity cDNA Reverse Transcriptase Kit (Applied Biosystems). Quantitative real-time PCR was performed using 1 µl of the cDNA preparation with SYBR-Green Reagents and a Vii7 qPCR analyzer (Applied Biosystems). Each DAF-16 target gene was amplified using PCR primers as described in (Senchuk *et al*., 2018) and compared to *act-3* (Key Resource Table).

## Statistical Analysis

Statistical analysis was carried out using GraphPad Prism software. For the analysis of two groups, a student t test was performed using two-tailed distribution for analysis involving two groups of samples. Fishers’ exact test was used for comparing groups of four. For each analysis, *P*<0.05 was considered as significant.

## ACKNOWLEDGEMENTS

We thank Ali Fazlollahi for technical assistance, Meera Sundaram (University of Pennsylvania) for comments and suggestions, Simon Wing, Richard Roy and Monique Zetka (McGill University) for reagents, Aimee Kao (UCSF) for *muIs113*, Patrick Hu (Vanderbilt University) for the *sgk-1* strains as well as Yanping Zhang, Wenhong Zhang and Meng-Qiu Dong (National Institute of Biological Sciences, Beijing) for generously sharing *daf-16(hq23)* ahead of publication. We thank Wormbase for information on genes and mutations. Some deletion mutations used in this study were provided by the National BioResource Project (Japan) and the International *C. elegans* Gene Knockout Consortium at the Oklahoma Medical Research Foundation, which is funded by the National Institutes of Health (NIH); and the University of British Columbia, which is funded by the Canadian Institute for Health Research (CIHR), Genome Canada, Genome B.C., the Michael Smith Foundation, and the NIH. Some strains were provided by the CGC, which is funded by NIH Office of Research Infrastructure Programs (P40 OD010440).

