## Supplemental Figures and Tables for "The Rab GTPase activating protein TBC-2 regulates endosomal localization of DAF-16 FOXO and lifespan"

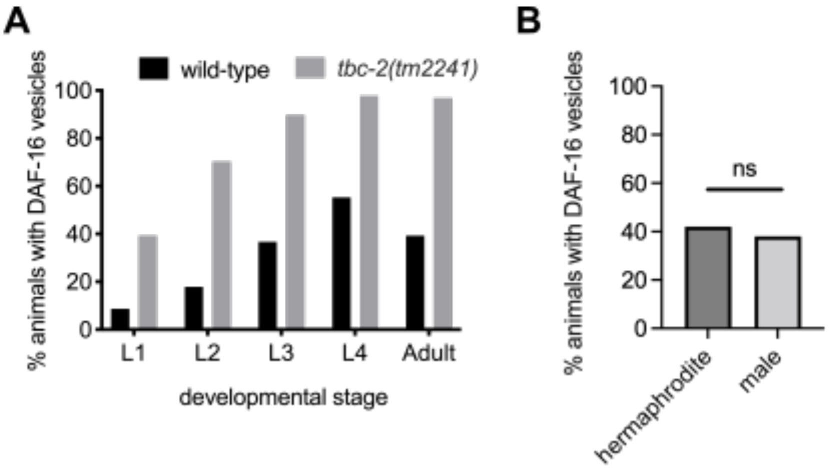


**Figure 1 – figure supplement 1. DAF-16::GFP localizes to vesicles in wild-type and *tbc-2* animals at all post-embryonic developmental stages and in wild-type males.**

(A) Bar graph of the percent animals with DAF-16a::GFP *(zIs356)* positive vesicles in the intestinal cells at larval stages L1-L4 and young adults of wild-type and *tbc-2(tm2241)* animals at 20°C. Fisher's exact test (graphpad.com) was used to determine that there is a significant increase the number of *tbc-2(tm2241)* animals with DAF-16a::GFP as compared to wild type at each developmental stage (L1: $P<0.05$, L2-adult: $P<0.0001$, $n$= 23 to 47 animals). (B) Bar graph of the percent *zIs356*/+ L4/young adult hermaphrodites ($n$=50) and males ($n$=47) with DAF-16a::GFP positive vesicles generated by crossing N2 males with QR779 *zIs356* hermaphrodites. ns, not significant.

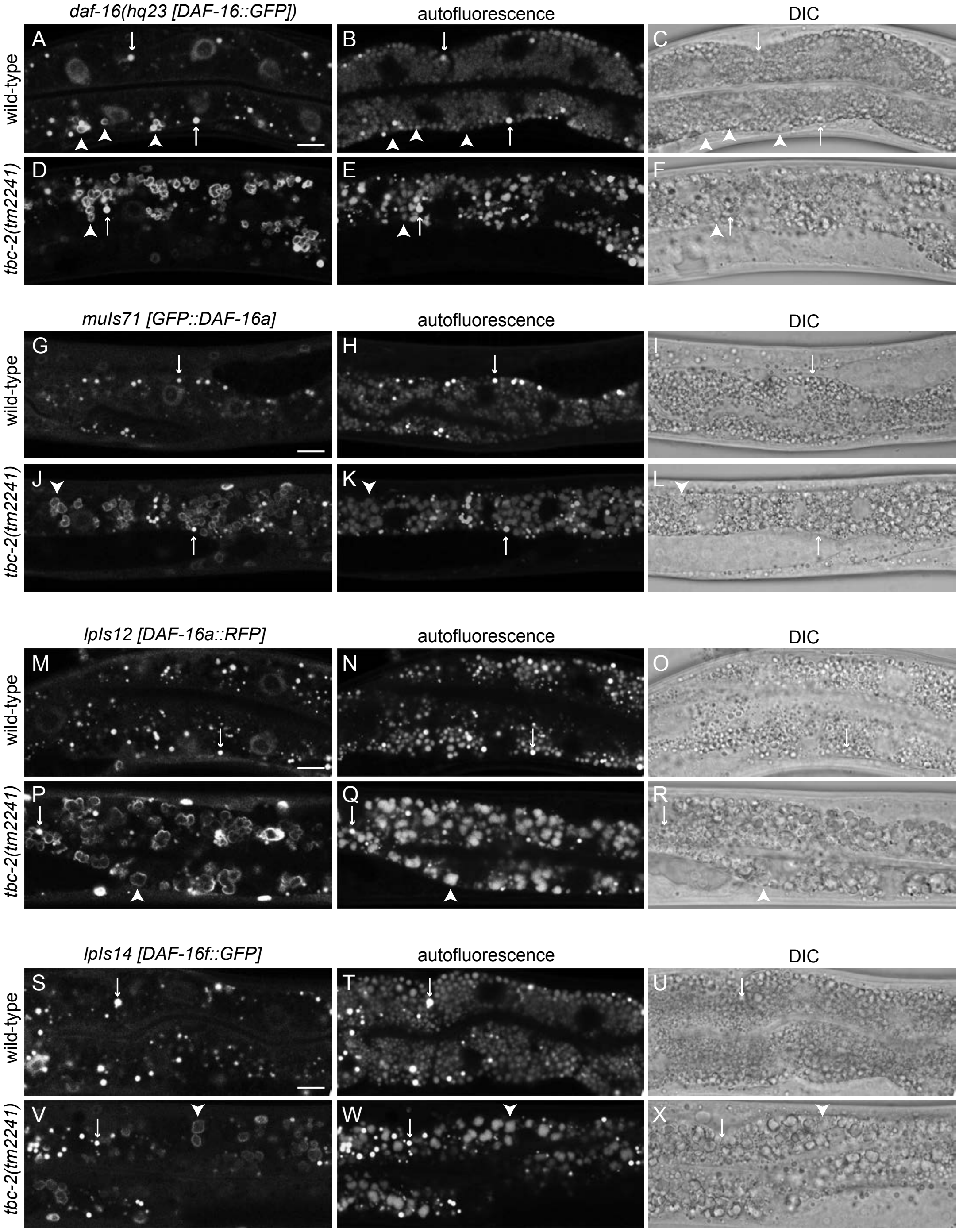
daf-16(hq23 [DAF-16::GFP])
autofluorescence
DIC
wild-type
tbc-2(tm2241)
A
B
C
D
E
F
muIs71 [GFP::DAF-16a]
autofluorescence
DIC
wild-type
tbc-2(tm2241)
G
H
I
J
K
L
lpIs12 [DAF-16a::RFP]
autofluorescence
DIC
wild-type
tbc-2(tm2241)
M
N
O
P
Q
R
lpIs14 [DAF-16f::GFP]
autofluorescence
DIC
wild-type
tbc-2(tm2241)
S
T
U
V
W
X

**Figure 1 – figure supplement 2. DAF-16 FOXO localizes to vesicles in intestinal cells**

Representative confocal and differential interference contrast (DIC) images of intestinal cells of wild-type (A-C, G-I, M-O and S-U) and *tbc-2(tm2241)* (D-F, J-L, P-R and V-X) animals expressing DAF-16::GFP *daf-16(hq23)* (A-F), *muIs71* GFP::DAF-16a (G-L), *lpIs12* DAF-16a::RFP (M-R) and *lpIs14* DAF-16f::GFP (S-X). Endogenously tagged DAF-16::GFP *daf-16(hq23)* is present on vesicles in both wild-type and *tbc-2(tm2241)* intestinal cells (A and D arrowheads). Arrows mark bright autofluorescent lysosome-related organelles that bleed through the GFP channel in these lower expressing strains. Vesicular localization of GFP::DAF-16a (G and J), DAF-16a::RFP (M and P) and DAF-16f::GFP (S and V) was only seen in *tbc-2(tm2241)* animals and not visible in wild-type backgrounds. Scale bars (A, G, M, S), 10µm.

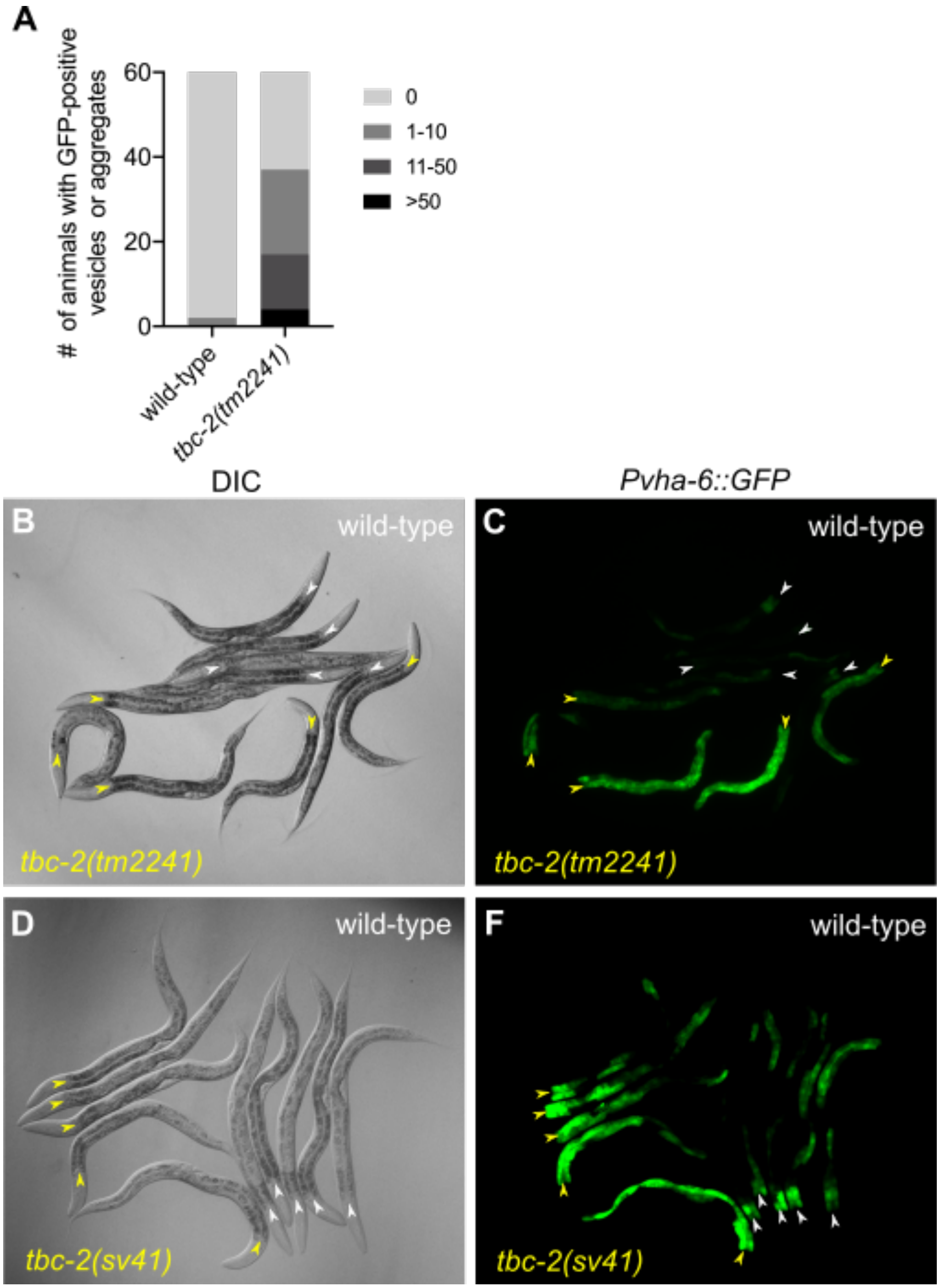


**Figure 1 – figure supplement 3. GFP localization and expression in wild-type and *tbc-2* mutants** (A) Grouped bar graph quantifying the number of wild-type and *tbc-2(tm2241)* L4 larvae with 0, 1-10, 11-50 or >50 GFP *(vhEx1[Pvha-6::GFP])* positive vesicles or aggregates (aggregates were included in the analyses which are not often seen with DAF-16a::GFP). (B-E)

DIC and epifluorescence images of wild-type and *tbc-2(tm2241)* (B,C) as well as wild-type and *tbc-2(sv41)* (D,E) animals expressing GFP under an intestine specific promoter, *vhEx1 [Pvha-6::GFP]*. White and yellow arrowheads mark the anterior of the intestine of wild-type and *tbc-2* mutant animals, respectively. Both *tbc-2* mutants have increased GFP expression as compared to wild-type animals. *tbc-2* mutants were distinguished from wild-type by the presence of enlarged vesicles using the 100X objective (not shown).

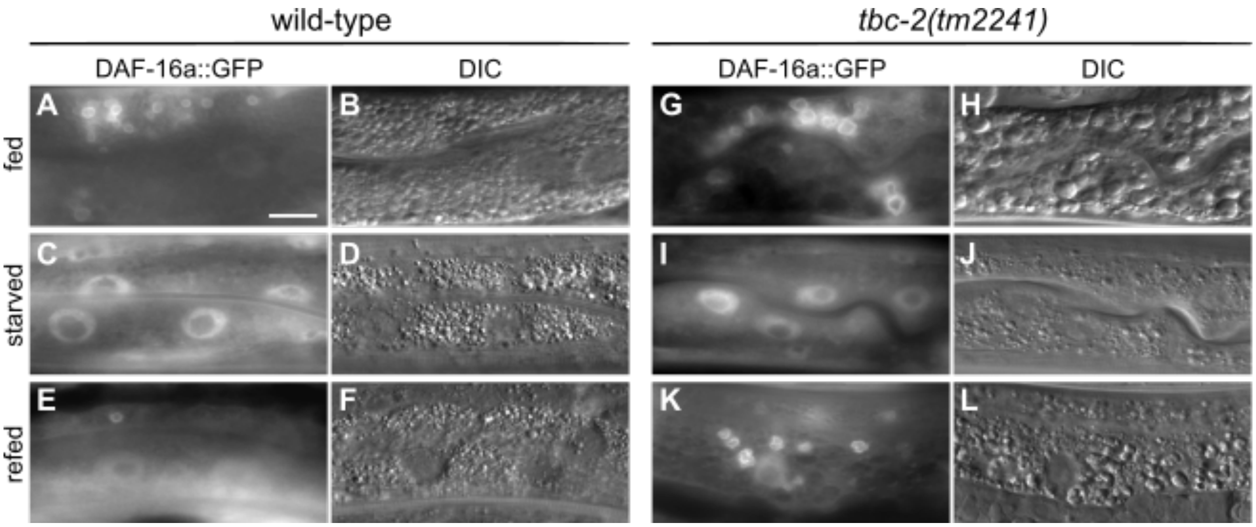


**Figure 4 – figure supplement 1. Endosomal DAF-16 is suppressed by acute starvation.** Epifluorescence (A,C,E,G,I,K) and DIC (B,D,F,H,J,L) images of wild-type (A-F) and *tbc-2(tm2241)* (G-L) animals under fed (A,B,G,H), 4-5 hours of starvation (C,D,I,J) and after 1-2 hours of refeeding (E,F.K.L). Scale bar (A), 10μm.

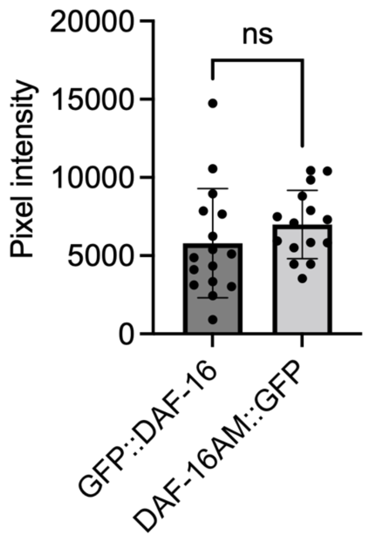


**Figure 4 – figure supplement 2. GFP::DAF-16 and DAF-16AM::GFP have similar expression levels** Bar graph depicting the mean pixel intensities of GFP fluorescence in the intestine of QR508 *tbc-2(tm2241); muIs71[GFP::DAF-16]* and QR697 *tbc-2(tm2241); muIs113[DAF-16AM::GFP]* animals. The difference was determined to be not significant (ns) in an unpaired t test.

| **Key Resources Table** | | | | |
|---|---|---|---|---|
| **Reagent type (species) of resource** | **Designation** | **Source or Reference** | **Identifier** | **Additional information** |
| Bacterial strain *(E. coli)* | HB101 | Caenorhabditis Genetics Center, CGC | | Nematode food source |
| Bacterial strain *(E. coli)* | HT115(DE3) | CGC | | RNAi feeding strain; Used in figures 3G,H and 4F,H |

| | | | | |
|---|---|---|---|---|
| Genetic reagent | empty vector (ev) L4440 RNAi feeding plasmid | Addgene | | Used in figures 3G,H and 4F,H |
| Genetic reagent | empty vector (EV) RNAi feeding strain | This study | | HT115(DE3) expressing L4440 RNAi clone, Used in figures 3G,H and 4F,H |
| Genetic reagent | RNAi feeding strains *rab-5* (I-4J01), *rab-7* (II-8G13), *akt-1* (V-7I17), *ftt-2* (X-5F07) and *par-5* (IV-6E06) | (Fraser *et al.*, 2000; Kamath *et al.*, 2003) | | Used in figures 3G,H and 4F,H |
| Nematode Strain (*C. elegans*) | CB1370 | CGC | *daf-2(e1370) III* | Used in figures 5 and 6 |
| Nematode Strain (*C. elegans*) | CF1407 | CGC | *daf-16(mu86) I; muIs71 [Pdaf-16a::GFP::daf-16a(bKO)) + rol-6(su1006)] X* | Used in figure 1 – figure supplement 2G-I |
| Nematode Strain (*C. elegans*) | HT1888 | CGC | *daf-16(mgDf50) I; unc-119(ed3) III; IpIs12[daf-16a::RFP + unc-119(+)]* | Used in figure 1 – figure supplement 2M-O |
| Nematode Strain (*C. elegans*) | HT1889 | CGC | *daf-16(mgDf50) I; unc-119(ed3) III; lpIs14 [daf-16f::GFP + unc-119(+)]* | Used in figure 1 – figure supplement 2S-U |
| Nematode Strain (*C. elegans*) | MQD1543 | CGC | *daf-16(hq23[DAF-16::GFP]) I* | Used in figure 1 – figure supplement 2A-C |
| Nematode Strain (*C. elegans*) | N2 | CGC | wild type | Used in figures 5, 6, and figure 1 – figure supplement 1B |
| Nematode Strain (*C. elegans*) | OH13908 | CGC | *daf-16(ot821[daf-16::mKate2::3xFlag]) I* | Used in figure 2B,D,F,H |
| Nematode Strain (*C. elegans*) | OH14125 | CGC | *daf-16(ot853[daf-16::linker::mNeonGreen::3xFlag::AID]) I* | Used in figure 2A,C,E,G,I |
| Nematode Strain (*C. elegans*) | QR15 | CGC | *tbc-2(tm2241) II* | Used in figures 5 and 6 |

| | | | | |
|---|---|---|---|---|
| Nematode Strain (*C. elegans*) | QR150 | This study | *tbc-2(tm2241) II; daf-2(e1370) III* | Used in figures 5 and 6 |
| Nematode Strain (*C. elegans*) | QR245 | (Skorobogata and Rocheleau, 2012) | *unc-119(ed3) III; vhEx1[Plin-31::GFP::rab-7 + Pvha-6::GFP + Cb-unc-119(+)]* | Used in figure 1 – figure supplement 3A-C |
| Nematode Strain (*C. elegans*) | QR272 | This study | *tbc-2(tm2241) II; zIs356 IV* | Used in figures 1B,D,F,H,K, 3G,H, 4A,D-F,H, figure 1 – figure supplement 1A, figure 4 – figure supplement 1A-F |
| Nematode Strain (*C. elegans*) | QR340 | This study | *zIs356 IV; pwIs480[Pvha-6::RFP::rab-5 + Cb unc-119(+)]* | Used in figure 3A-C |
| Nematode Strain (*C. elegans*) | QR505 | This study | *tbc-2(tm2241) II; lpIs14 [daf-16f::GFP + unc-119(+)]* | Used in figure 1 – figure supplement 2V-X |
| Nematode Strain (*C. elegans*) | QR508 | This study | *tbc-2(tm2241) II; muIs71 [Pdaf-16a::GFP::daf-16a(bKO)) + rol-6(su1006)] X* | Used in figure 4I, figure 4 – figure supplement 2 |
| Nematode Strain (*C. elegans*) | QR646 | This study | *daf-18(e1375) zIs356 IV* | Used in figure 4D |
| Nematode Strain (*C. elegans*) | QR647 | This study | *daf-18(e1375) zIs356 IV* | Used in figure 4D |
| Nematode Strain (*C. elegans*) | QR648 | This study | *daf-18(e1375) zIs356 IV* | Used in figure 4D |
| Nematode Strain (*C. elegans*) | QR651 | This study | *zIs356 IV; pwIs429 [Pvha-6::mCherry::rab-7 + Cb unc-119(+)]* | Used in figure 3D-F |
| Nematode Strain (*C. elegans*) | QR655 | This study | *tbc-2(tm2241) II; vhEx1* | Used in figure 1 – figure supplement 3A-C |
| Nematode Strain (*C. elegans*) | QR658 | This study | *daf-18(ok480) zIs356 IV* | Used in figure 4D |

| | | | | |
|---|---|---|---|---|
| Nematode Strain (*C. elegans*) | QR659 | This study | *daf-18(ok480) zIs356 IV* | Used in figure 4D |
| Nematode Strain (*C. elegans*) | QR660 | This study | *daf-18(ok480) zIs356 IV* | Used in figure 4D |
| Nematode Strain (*C. elegans*) | QR661 | This study | *daf-18(ok480) zIs356 IV* | Used in figure 4D |
| Nematode Strain (*C. elegans*) | QR662 | This study | *daf-18(ok480) zIs356 IV* | Used in figure 4D |
| Nematode Strain (*C. elegans*) | QR664 | This study | *tbc-2(tm2241) II; lpIs12 [daf-16a::RFP + unc-119(+)]* | Used in figure 1 – figure supplement 2P-R |
| Nematode Strain (*C. elegans*) | QR688 | This study | *zIs356 IV; akt-2(ok393) X* | Used in figure 4E |
| Nematode Strain (*C. elegans*) | QR689 | This study | *zIs356 IV; akt-2(ok393) X* | Used in figure 4E |
| Nematode Strain (*C. elegans*) | QR697 | This study | *tbc-2(tm2241) II; muIs113 [Pdaf-16::daf-16AM::gfp + rol-6(su1006)]* | Used in figure 4I, figure 4 – figure supplement 2 |
| Nematode Strain (*C. elegans*) | QR729 | This study | *daf-16(hq23[DAF-16::GFP]) I; tbc-2(tm2241) II* | Used in figure 1 - figure supplement 2D-F |
| Nematode Strain (*C. elegans*) | QR779 | This study | *zIs356[Pdaf-16::daf-16a/b(D484V)::GFP + rol-6(su1006)] IV* | TJ356 outcrossed to N2 6; used in figures 1I, figure 1 – figure supplement 1B and 4C,G |
| Nematode Strain (*C. elegans*) | QR807 | This study | *tbc-2(tm2241) II; zIs356 IV* | Derived from QR779; used in figure 1J |
| Nematode Strain (*C. elegans*) | QR851 | This study | *daf-2(e1370) III; zIs356 IV* | Used in figure 4C |
| Nematode Strain (*C. elegans*) | QR852 | This study | *daf-2(e1370) III; zIs356 IV* | Used in figure 4C |
| Nematode Strain (*C. elegans*) | QR853 | This study | *daf-2(e1370) III; zIs356 IV* | Used in figure 4C |

| | | | | |
|---|---|---|---|---|
| Nematode Strain (*C. elegans*) | QR869 | This study | *tbc-2(sv41) II; daf-2(e1370) III* | Used in figures 5 and 6 |
| Nematode Strain (*C. elegans*) | QR910 | This study | *vhEx1 [Plin-31::GFP::rab-7 + Pvha-6::GFP + Cb-unc-119(+)]* | Used in figure 1 – figure supplement 2D,E |
| Nematode Strain (*C. elegans*) | QR915 | This study | *tbc-2(sv41) II; vhEx1* | Used in figure 1 – figure supplement 2D,E |
| Nematode Strain (*C. elegans*) | QR1057 | This study | *daf-16(ot853[daf-16::linker::mNeonGreen::3xFlag::AID]) I; tbc-2(tm2241) II* | Used in figure 2I |
| Nematode Strain (*C. elegans*) | TJ356 | CGC | *zIs356[Pdaf-16::daf-16a/b(D484V)::GFP + rol-6(su1006)] IV* | Used in figures 1A,C,E,G, 3G,H, 4A,D-F,H, figure 1 – figure supplement 1A, figure 4 – figure supplement 1A-F |
| Nematode Strain (*C. elegans*) | UP1224 | (Chotard et al., 2010a) | *tbc-2(sv41) II* | Used in figures 5 and 6 |
| Nematode Strain (*C. elegans*) | | (Chen *et al.*, 2013) | *daf-16(mu86) I; zIs356 IV; sgk-1(ft15) X* | Used in figure 4G |
| Nematode Strain (*C. elegans*) | | (Chen *et al.*, 2013) | *daf-16(mu86) I; zIs356 IV; sgk-1(ok538) X* | Used in figure 4G |
| Nematode Strain (*C. elegans*) | | This study | *zIs356 IV; him-5(e1467) V* | Used in figure 1 – figure supplement 1B |
| Sequence based reagent | *act-3* qRT-PCR primers | (Senchuk *et al.*, 2018) | Forward 5'- TGC GAC ATT GAT ATC CGT AAG G -3'<br>Reverse 5'- GGT GGT TCC TCC GGA AAG AA -3' | Used in figure 6 |
| Sequence based reagent | *sod-3* qRT-PCR primers | (Senchuk *et al.*, 2018) | Forward 5'- AAA GGA GCT GAT GGA CAC TAT TAA GC -3'<br>Reverse 5'- AAG TTA TCC AGG GAA CCG AAG TC -3' | Used in figure 6 |
| Sequence based reagent | *dod-3* qRT-PCR primers | (Senchuk *et al.*, 2018) | Forward 5'- AAG TGC TCC GAT TGT TAC GC -3'<br>Reverse 5'- ACA TGA ACA CCG GCT CAT TC -3' | Used in figure 6 |

| | | | | |
|---|---|---|---|---|
| Sequence based reagent | *mtl-1* qRT-PCR primers | (Senchuk *et al.*, 2018) | Forward 5'- ATG GCT TGC AAG TGT GAC TG -3'<br>Reverse 5'- GCT TCT GCT CTG CAC AAT GA -3' | Used in figure 6 |
| Sequence based reagent | *ftn-1* qRT-PCR primers | (Senchuk *et al.*, 2018) | Forward 5'- GAG TGG GGA ACT GTC CTT GA -3'<br>Reverse 5'- CGA ATG TAC CTG CTC TTC CA -3' | Used in figure 6 |
| Sequence based reagent | *gpd-2* qRT-PCR primers | (Senchuk *et al.*, 2018) | Forward 5'- CTC CAT CGA CTA CAT GGT CTA CTT G -3'<br>Reverse 5'- AGC TGG GTC TCT TGA GTT GTA GAC -3' | Used in figure 6 |
| Sequence based reagent | *icl-1* qRT-PCR primers | (Senchuk *et al.*, 2018) | Forward 5'- TGT GAA GCC GAG GAC TAC CT -3'<br>Reverse 5'- TCT CCG ATC CAA GCT GAT CT -3' | Used in figure 6 |
